# Mechanical Heterogeneity in the Bone Microenvironment as Characterised by Atomic Force Microscopy

**DOI:** 10.1101/2020.02.25.964791

**Authors:** X. Chen, R. Hughes, N. Mullin, R. J. Hawkins, I. Holen, N. J. Brown, J. K. Hobbs

## Abstract

Bones are structurally heterogeneous organs with diverse functions that undergo mechanical stimuli across multiple length scales. Mechanical characterisation of the bone microenvironment is important for understanding how bones function in health and disease. Here we describe the mechanical architecture of cortical bone, the growth plate, metaphysis and marrow in fresh murine bones, probed using atomic force microscopy in physiological buffer. Both elastic and viscoelastic properties are found to be highly heterogeneous with moduli ranging over 3 to 5 orders of magnitude, both within and across regions. All regions include extremely soft areas, with moduli of a few Pascal and viscosities as low as tens Pa⋅s. Aging impacts the viscoelasticity of the bone marrow strongly but has limited effect on the other regions studied. Our approach provides the opportunity to explore the mechanical properties of complex tissues at the length scale relevant to cellular processes and how these impact on aging and disease.

**SIGNIFICANCE:** The mechanical properties of biological materials at cellular scale are involved in guiding cell fate. However, there is a critical gap in our knowledge of such properties in complex tissues. The physiochemical environment surrounding the cells in *in-vitro* studies differs significantly from that found *in vivo*. Existing mechanical characterisation of real tissues are largely limited to properties at larger scales, structurally simple (*e.g.* epithelial monolayers) or non-intact (*e.g.* through fixation) tissues. In this paper, we address this critical gap and present the micro-mechanical properties of the relatively intact bone microenvironment. The measured Young’s moduli and viscosity provide a sound guidance in bioengineering designs. The striking heterogeneity at supracellular scale reveals the potential contribution of the mechanical properties in guiding cell behaviour.

## INTRODUCTION

It is increasingly clear that mechanical properties and forces play an essential role in controlling many aspects of cell biology of tissues and organs including growth, migration, differentiation, homeostasis and communication (1–3). How these fundamental cellular processes are influenced by such forces requires an understanding of the mechanical properties at a subcellular scale but in the context of the tissue, organ, or even the whole organism. Obtaining this information is particularly challenging as most traditional methods for tissue processing (*e.g*. sectioning) or fixation radically change the mechanical properties of at least part of the structure (*e.g*. by freezing and thawing). Similarly, obtaining the relevant spatial and temporal resolution with sufficient force sensitivity to systematically determine the relevant mechanical variation at the different cell and tissue scales requires new specialised approaches. Here we present an atomic force microscopy (AFM) based approach for tackling these issues allowing the study of bones, a unique and mechanically complex organ.

Bone tissue consists predominantly of two types of bone, cortical and trabecular. The cortical bone (also known as dense bone) is structurally compact and bears the load of the body’s weight. The trabecular bone (also known as cancellous bone or spongy bone) has a more loosely organized structure and highly metabolic activity (4). Both types of bone contain inorganic (bone mineral and water) and organic components (bone cells and matrix) that form complex, interconnected structures (5). In addition to being central to mechanical support, bone contains bone marrow which plays a significant role in mammalian physiology through hematopoiesis and by regulating immune and stromal cell trafficking (6). Bone tissues undergo continuous dynamic remodelling in response to changes in mechanical loading and during the homeostatic repair of microdamage occurring through normal ‘wear and tear’. The balance between bone deposition and resorption is orchestrated by bone cells, called osteoblasts and osteoclasts, respectively (4, 5). Osteoblasts are derived from mesenchymal stem cells (MSCs) and osteoclasts are derived from hematopoietic stems cells (HSCs), both of which are resident in bone marrow (6). In addition there are a multitude of progenitor cells, immune cells, stromal cells and adipocytes involved in maintaining the homeostasis of bone.

The mechanical properties of bone are fundamental to the function of the skeleton (7) and changes in bone loading can influence bone turnover, a process whereby bone formation and absorption is in physiological balance, thereby preventing the net gain or loss of bone tissue (8). Investigations of bone mechanics are necessary to fully understand how the mechanical properties of the bone microenvironment (BMev) influence the biological functions such as bone turnover, and have been well developed in recent decades. *In-vitro* models mimicking the BMev are commonly used to study the role of mechanical properties in bone function and related diseases, such as cancer-induced bone metastasis (9, 10). However, these *in vitro* models do not fully recapitulate the complexity of the *in vivo* BMev and hence lack fundamental components of the underlying biology (10).

Numerous studies using a variety of techniques have characterised the mechanical properties of bone at multiple length scales. Elastic and viscoelastic studies on both cortical and trabecular bones generally give Young’s/shear moduli in the GPa range, though the results vary due to different experimental conditions (11–22). The Young’s/shear modulus of trabecular bone is slightly lower than that of the cortical bone and occasionally as low as 10s - 100s MPa (23, 24). The growth plate is more compliant than both cortical and trabecular bone tissues, with the Young’s modulus ranging from 300 kPa to 50 MPa (25–28). In contrast, most studies on bone marrow have considered it as a purely viscous tissue (29), reporting viscosity values ranging from below 1 Pa⋅s to 100 Pa⋅s (29–31).

The structural complexity of bone samples can hinder accurate mechanical monitoring. A number of approaches have been taken to overcome this, through exposing bone to freezing (32, 33), dehydration (34), jet washing (35), polishing (12, 19, 36) and homogenising (37, 38) in addition to sectioning or fracturing. Although differences in the mechanical properties of the BMevs may be extracted from samples prepared via these methods, such treatments substantially modify the surface structures (35) and consequently cause a shift in the measured mechanical properties (14, 19). However, the ability to quantify the mechanical properties of tissues at different length scales is critical in building models for theoretical simulations or in tissue engineering to replicate key features of such tissues. A recent study on intact bone marrow demonstrated both the feasibility and advantages of mechanical measurements using minimally deconstructed samples (39). In contrast to other studies, this work identified that the bone marrow is predominantly elastic, and suggested that the extracellular matrix contribution to the mechanical properties from fresh samples is essential to explain the observed heterogeneity. The accessible scale in this study was not sufficient to reveal the mechanical heterogeneity of bone, and suggested that techniques suitable for smaller scale characterisation are necessary.

Several methods to evaluate the local (*i.e.* cellular or subcellular) mechanical properties of biological tissues have emerged over the last decade, including optical (40, 41) and acoustic (42) techniques. However, the structural and mechanical heterogeneity of the BMev makes the application of these approaches particularly challenging. AFM enables characterisation at length scales ranging from nm to tens of μm (25, 43–45), which is of particular relevance for understanding the biological mechanisms arising at the molecular and cellular level.

Here we focused on measuring the subcellular mechanical properties of the non-load bearing components of untreated bone on the internal surfaces (except for longitudinal splitting, which is necessary to access the tissue). Mechanical measurements were made using colloidal probe AFM, as opposed to nanoindentation (19, 34) or AFM with conventional sharp probes (13, 14, 17, 25), due to the extreme softness of non-mineralised components in the BMev. Point measurements were performed randomly over a macro-scale (> 100 µm) of bone surfaces. Both the elastic and viscoelastic properties of different regions of the BMev were quantified by fitting data obtained from force-distance and creep (*i.e.* strain relaxation) curves acquired with the AFM with appropriate mechanical models. Bones from young (6-8 weeks) and mature (11-13 weeks) mice were compared to determine any mechanical changes due to age-associated skeletal maturation. Finally, AFM force mapping was used to collect images (max. size 100 × 100 µm^2^) that represented the heterogeneity of the BMev morphology and mechanics at a supracellular scale (from 5 µm to 100 µm). The implications of our findings for the potential correlation with biological functions in bones are also discussed.

## MATERIALS AND METHODS

### Animals

All experiments involving animals were approved by the University of Sheffield project Applications and Amendments (Ethics) Committee and conducted in accordance with UK Home Office Regulations, under project license number 70/8964 (NJB). Mice were housed in a controlled environment with 12 hours light/dark cycle, at 22°C. Mice had access to food and water *ad libitum*, Teklad Diet 2918 (Envigo, UK). A total of 21 mice were used in these studies.

### Bone sample preparation

Femurs from both young (6 to 8 weeks old, 9 bones from 8 mice) and mature (11 to 13 weeks old, 24 bones from 13 mice) mice were used in this study unless otherwise stated. The mice were culled by cervical dislocation and the hind limbs detached from the pelvis at the femoral head. Muscle tissue connected to the bone was removed and the femur separated from the tibia at the knee joint. The dissected femurs were placed in phosphate buffered saline (PBS) (Lonza, US) at 4°C and split prior to measurement, which was within 36h of culling. A custom designed tool with a razor blade (Fig. S1a) was used to longitudinally fracture the bone, which was immobilised on a platform using a two component dental impression putty (Provil Novo Light, Kulzer, UK). The split bone was then immersed in PBS and immobilised in a petri-dish (TPP, Switzerland) with the same dental impression material. Care was taken to maintain hydration of the exposed bone surface during the entire process.

### Colloidal probe cantilever preparation

Rectangular Si_3_N_4_ cantilevers with a nominal spring constant of 0.02 N/m (MLCT cantilever B, Bruker, USA) were used for all AFM measurements. A polystyrene microsphere with 25 µm diameter (Sigma Aldrich, USA) was attached to the cantilever using a UV-curing adhesive (NOA 81, Norland Inc., USA), using an AFM (Nanowizard III, JPK, Germany) and combined with an inverted optical microscope (Eclipse Ti, Nikon, Japan) as a micro-manipulator. Before attachment of the colloidal probe, the spring constant of each cantilevers was determined in air using the thermal noise method (46). To prevent strong non-specific binding between the probe and sample surface, the cantilever with attached colloidal probe and cantilever holder were passivated by 20 min immersion in 10 mg/mL bovine serum albumin (BSA), (Sigma Aldrich, USA). Excess BSA was removed by washing with 5 × 1 mL PBS.

### AFM force spectroscopy and mapping

AFM measurements were all performed on a Nanowizard III system (JPK, Germany) using a heated sample stage (35 to 37 °C). The cantilever deflection sensitivity was calibrated prior to each measurement, from the hard contact regime of force-distance curves (average of 10 repeated curves) obtained from a clean petri-dish containing PBS. Prepared bone samples in PBS (Fig. 1a) were then mounted on the sample stage and probed after the temperature of the liquid had stabilised. Force-distance curves (Fig. 1b) were acquired at randomly selected positions from different regions of interest towards the distal end of each femur, as shown in Fig. 1a. For the majority of samples, 5 to 10 positions within each BMev region were measured from each femur, but on occasion the number varied beyond this range depending on sample quality (*e.g.* surface roughness). The approach speed was 1 µm/s unless otherwise stated. Subsequently, curves with 3 s dwell under constant force (0.5 nN), *i.e.* creep curves, were also acquired from the same position (Fig. 1d). For both force-distance and creep curves, a minimum of 3 measurements were taken at each location.

**Figure 1.**
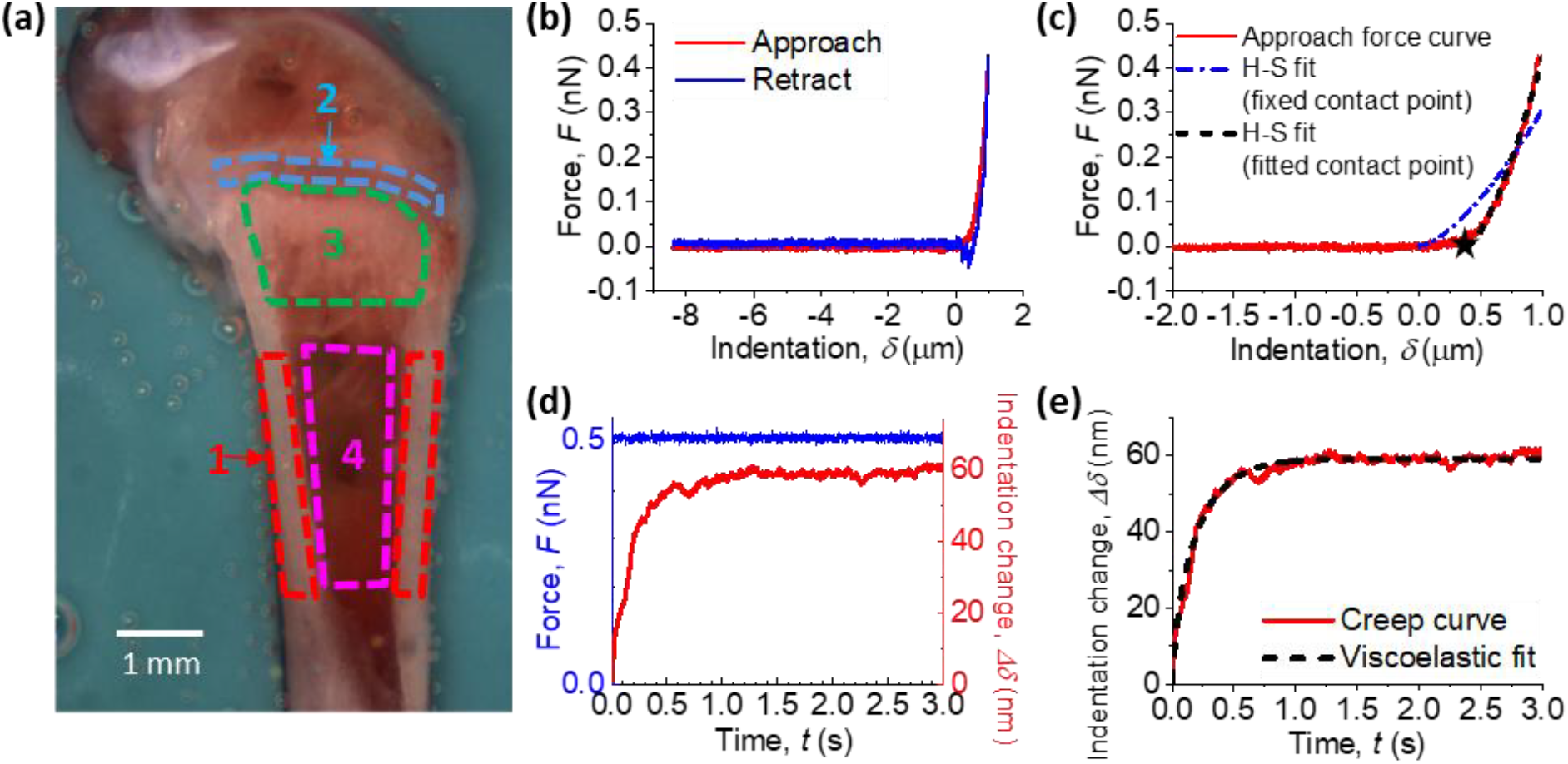
Methods of measuring the mechanical properties on internal bone surfaces (coloured version available online). (a) Bright field image of a prepared bone surface for AFM characterisation Four regions of interest are indicated by coloured dashed lines: (1) cortical bone (*red*), (2) the growth plate (*blue*), (3) the metaphysis (*green*) and (4) bone marrow in diaphysis (*magenta*). (b) Representative force (*F*) vs. indentation (*δ*) curve obtained during AFM probe approach (*red*) and retract (*blue*) taken on the bone surface. The contact position (δ = 0) was determined from the approach curve using triangle thresholding method described in the Supporting Material. (c) Examples of Hertz-Sneddon (H-S) model fits to the approach segment of a force-indentation curve (*red*) H-S fit either using a fixed contact point (*blue dash-dotted line*) determined using the triangle thresholding method (described in the Supporting Material), or with the contact point as a free fitting parameter (*black dashed line*). The star (★) shows the position of the contact point determined by the fit. (d) Representative creep curve (*Δδ vs. t*) obtained from the dwell segment of a force curve taken on the bone surface. The applied force (*blue*) was held constant for 3 s while the indentation depth increased (*red*) due to the material being viscoelastic rather than purely elastic. (e) Example of the viscoelastic model fit (*black dashed line*) on a creep curve (*red line*).

A two-dimensional array of force-distance curves (*i.e.* an AFM map, one force curve per pixel) was then produced at randomly selected areas in the different bone regions, with 30 µm/s approach speed. The faster approach speed for force maps was chosen to minimise data collection time and consequent degradation of the sample. Before obtaining maps for analysis (typically 1-1.6 µm per pixel), low resolution “survey” maps (10 to 15 µm per pixel) were collected to ensure that the major features in the selected region could be mapped without exceeding the 15 µm Z range of the AFM.

All AFM measurements (force-distance curves, creep curves and force maps) on a single bone sample could typically be collected in 2 to 6 h. The measured bone surfaces were stable during this time (*i.e.* there was no significant dissociation or significantly different features in force curves observed). Reference force-distance curves were acquired on a petri-dish at the end of the measurements to check for colloidal probe contamination, with no significant contaminants (expected to manifest as strong adhesive features in the retract segment of the curves) found.

Raw data were exported as .txt format using JPK Data Processing and imported into customised algorithms in MATLAB (generated in-house by XC) for all subsequent analyses. Data processing scripts are available upon request.

### Processing and analysis of force-distance curves

Raw force-distance curves were first converted into force (*F*) - indentation (*δ*) curves (Fig. 1b) by applying baseline correction, determining the contact point and subtracting the cantilever deflection from the Z displacement in the contact regime (see Supporting Methods and Fig. S1b-e). Curves were fitted to a Hertz-Sneddon model (47) using a custom Matlab routine, with the Young’s modulus (*E*_*H-S*_) and a virtual contact point as free fitting parameters (see Supporting Methods). The *E*_*H-S*_ at any measured position was determined by the mean value resulting from fitting to all repeated *F*-*δ* curves at the same position, unless otherwise specified. The quality of *F*-*δ* curves (*i.e.* tilt in baselines) and the fitting quality variation only had a minor impact on the results, as discussed in the Supporting Methods. It is important to note that the analysis described in this section implicitly assumes that the mechanical response of the BMev is purely elastic.

### Processing and analysis of creep curves

Raw force (*F*) – time (*t*) curves from the dwell segment of force-distance curves were checked and discarded if the force could not be maintained constant by the feedback loop. The Z displacement value at the start of the dwell segment was subtracted from the displacement-time curve to yield an indentation variation (*Δδ*) - time (*t*) curve (creep curve). This was fitted to both a Standard Linear Solid (SLS) model (Fig. S1f) and a Kelvin-Voigt (K-V) model (Fig. S1g), based on the theory developed by Cheng *et. al.* (48) (see Supporting Methods). The instantaneous elastic modulus (*E*_*1*_), delayed elastic modulus (*E*_*2*_) and viscosity (*η*) were free fitting parameters using the SLS model, and the Young’s modulus (*E*_*K-V*_) and viscosity (*η*) using the K-V model. Results were obtained from the mean value of repeated measurements at each position and those with low fit quality (*R*^2^ < 0.9) were discarded if not specified.

### Processing of AFM map data

Three types of maps that graphically represented topography and mechanical properties of the scanned areas were generated from the AFM maps. Firstly, the trigger point map was constructed from a vertical position where the interaction force between the probe and the sample reached the trigger value. This represents the topography of the surface, including the indentation of the probe into the sample. Secondly, the contact point map was constructed from the vertical position where the probe was deemed to have made initial contact with the sample surface, determined from the contact point found for every force curve in the map using the triangle thresholding method, as shown in Fig. S1b. This represents an approximation to the topography at zero load and indentation. Thirdly, the elastic map was constructed from the Young’s modulus *E*_*H-S*_ extracted from all force curves in the map using the Hertz-Sneddon model with a fitted contact point. For both the contact point map and the mechanical map, pixels with curves that could not be properly fitted due to not having a clear non-contact baseline, contact point and monotonic contact regime were considered as “missing data” and coloured black in the figures.

### Statistical analysis

Statistical analyses were performed using OriginPro software. A normality test was applied to all distributions prior to any further analysis. Data were then analysed by the Kruskal-Wallis test for comparison between different groups. A statistically significant difference was defined as *p* < 0.05.

## RESULTS AND DISCUSSION

Four regions of interest (Fig. 1a) within the BMev were selected on the surface of split bones for AFM characterisation, due to their diverse composition and function: (i) cortical bone, (ii) growth plate, (iii) metaphysis and (iv) bone marrow in the diaphysis. The four regions of interest are shown by coloured outlines in the bright field image (Fig. 1a). In long bones such as the femur, the growth plate (also known as the epiphyseal plate or physis) is found at each end. This is where longitudinal bone growth takes place and is composed of a large number of chondrocytes and proteinaceous matrix. The narrow portion located just below the growth plate, the metaphysis (4), was also examined. Previous preclinical *in vivo* studies from our group have identified this as the predominant region where circulating metastatic cancer cells home to and colonise in bone (49, 50). In the metaphysis, measurements were made on either the trabecular bone tissue or adjacent soft tissue in the surrounding marrow. In the diaphysis, we measured both the cortical bone, which comprises the outer shaft of the long bone, and the central bone marrow cavity.

Only bones with an acceptably smooth surface topography after splitting were used for AFM measurements. However, even on these specimens, the height range of the surface profile could still vary by several hundred micrometres across the whole bone. Tilt or curvature of the split surface occasionally caused fouling of the cantilever, or cantilever holder on parts of the bone that were higher than the region of interest. In these cases it was not possible to measure all regions of interest on the same specimen.

### Point force-indentation measurements reveal mechanical heterogeneity spanning orders of magnitude over each of the different regions of the bone microenvironment, with all regions containing very soft (1-10 Pa) areas

To probe the overall mechanical profile for each bone region, we first applied point force (*F*) -indentation (*δ*) measurements at randomly selected positions within the specified region. A representative *F*-*δ* curve acquired on the surface of a split bone is shown in Fig. 1b (see Supporting Material and Fig. S1b-e for details of conversion of raw deflection-distance curves to *F*-*δ* curves). In the majority of curves, features associated with plastic deformation (yield points, plateaus etc.) were not observed, indicating that elastic or viscoelastic analyses were appropriate. For simplicity, the Young’s modulus *E*_*H-S*_, together with a virtual contact point, was obtained from Hertz-Sneddon model fits to the *F*-*δ* curves (Fig. 1c) assuming a purely elastic response. Data include bones from both young and mature mice.

Histograms of *E*_*H-S*_ for each bone region are shown in Fig. 2 (note the logarithmic scale of the horizontal axes), and follow neither normal nor log-normal distributions. The median *E*_*H-S*_ is (i) cortical bone: 0.29 kPa, (ii) growth plate: 91 Pa, (iii) metaphysis: 17 Pa and (iv) bone marrow: 6.7 Pa. The mean *E*_*H-S*_ is (i) cortical bone: 2.7 kPa, (ii) growth plate: 0.19 kPa, (iii) metaphysis: 0.42 kPa and (iv) bone marrow: 0.14 kPa. These data indicate that the BMev is very soft.

**Figure 2.**
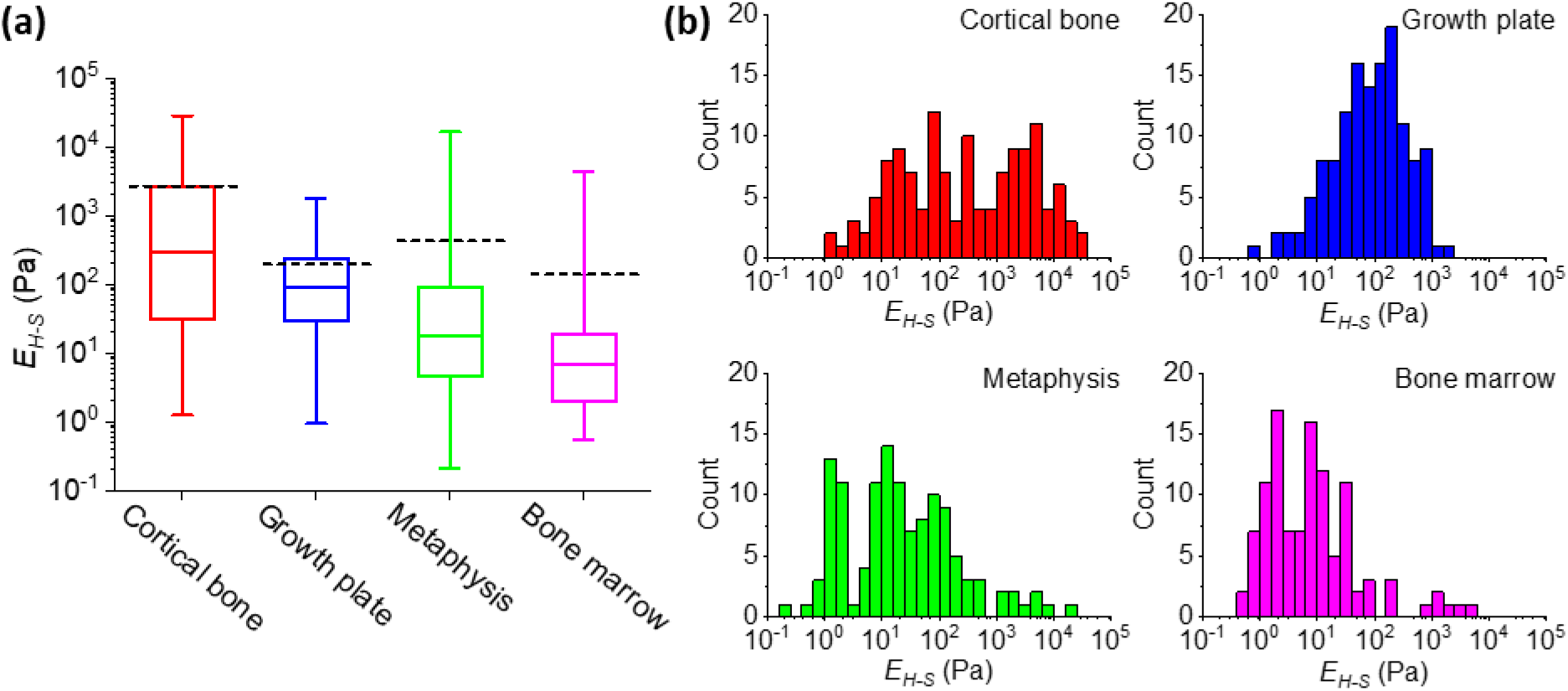
Elastic properties of different bone regions (coloured version available online). (a) The Young’s modulus (*E*_*H-S*_) of different regions in the BMev calculated using the Hertz-Sneddon model with fitted contact point. The central box spans the lower quartile to the upper quartile of the data. The solid line inside the box shows the median and whiskers present the lower and upper extremes. The mean values are indicated by black dashed lines. Data were obtained at randomly selected positions within regions of interest on bones from both young and mature mice. The Young’s modulus reported at each position is the mean value of individual fits to all force-indentation curves taken at that location. Curves with strongly tilted baselines (*i.e*. *d*_*b*_ > *d*_*c*_/4, as in Fig. S1c) were discarded. Results from low quality fittings (*i.e*. R^2^ < 0.9) were also discarded. (b) Histograms of the *E*_*H-S*_ of different bone regions, as in (a). The corresponding histograms in linear scale are presented in Fig. S2.

The distributions of *E*_*H-S*_ are very broad in all regions of interest, covering several orders of magnitude (see Fig. S2 for data on a linear scale), revealing the extreme mechanical heterogeneity within each region, which is consistent with the known heterogeneous structure of bone (12, 26, 33).

The mechanical distributions vary significantly between different bone regions (*p* < 10^−4^ under Kruskal-Wallis test). Histograms of *E*_*H-S*_ measured for cortical bone show a wide distribution that lacks a sharp peak with a large portion of data at relatively high *E*_*H-S*_ (*i.e. E*_*H-S*_ > 10^3^ Pa), compared to the mechanical distribution of the other bone regions measured. This is in agreement with the high level of mineralisation in cortical bone (5). The mechanical distribution of both the growth plate and bone marrow are narrower, reflecting the more homogeneous composition and/or structure compared to other bone regions (51). The measured modulus of the metaphysis covers the widest range, due to the presence of both bone tissue and bone marrow in this region, enhancing the structural and consequently mechanical heterogeneity.

Our approach has limits in acquiring data beyond the range shown in the histograms. At the low stiffness end of the distribution (Pa), errors are dominated by inaccuracies in determining the contact point, and tend to lead to an overestimate of the extracted moduli, so the distributions may extend to still lower moduli. At the high stiffness end (100s kPa) there is insufficient indentation at the maximum (trigger) forces used which may lead to an underestimate in the extracted moduli. As a control, the *E*_*H-S*_ extracted from hard wall *F*-*δ* curves, from the polystyrene petri-dish (expected *E*_*H-S*_ ~ 3 GPa) under the same experimental conditions, does not exceed 10s to 100s kPa, which is the upper limit of *E*_*H-S*_ using the current AFM setup (that is focused on measuring soft components). The experimental setup is not sensitive to the GPa moduli of calcified cortical and trabecular bone previously described (11–15), as this is beyond the scope of this study.

However, our approach has the important advantage of being able to measure the subcellular properties of soft tissues, i.e. the cellular microenvironment. The Young’s moduli found in the current study are orders of magnitude lower than the values reported in the majority of previous studies (11–28, 34). Direct comparison is difficult, but the ability to measure at the length scale of the cellular components is likely to contribute to the difference. Previous experiments performed with non-AFM indentation approaches (19, 34) normally average over large areas (hundreds μm^2^) and thus are unable to determine the properties of the smaller scale. Meanwhile, the low spring constant cantilever and relatively large contact radius probe were selected for use in the current study after extensive preliminary experiments, and provide sufficient force sensitivity to assess subcellular scale mechanics in soft tissue. This is in contrast to using conventional sharp AFM probes and enables accurate measurements of the soft components in the BMev, rather than simply being able to obtain measurements from the calcified regions (11–15).

In previous studies mechanical characterisation of the bone marrow is reported to be distinct from other regions within the BMev. The marrow region is traditionally considered as a viscous fluid (29) and there are only a few studies describing the elastic properties.. However, a recent study suggested that the bone marrow is predominantly elastic, and reported an effective Young’s modulus ranging from 0.1 to 10.9 kPa measured at physiological temperature (39). This value obtained from porcine bone is similar to that obtained in the current study from murine bone, the difference in the resultant moduli most likely due to the use of rheological measurements instead of indentation. It is important to note that the bone marrow used both in the previous (36) and the current study, is essentially intact with no processing (such as fixation) prior to measurement of the mechanical properties.

The H-S model with fitted contact point does not fit the entire *F*-*δ* curve perfectly (Fig. 1c). This is likely because the H-S model assumes that the probed material is homogeneous and linearly elastic, whereas the BMev is known to be structurally heterogeneous (5) (Additional errors in the model are discussed in the Supporting Materials). Elastic models describing discontinuous materials (*e.g.* cell-polymer brush model (52)) may be helpful for improved determination of the elasticity of the BMev. Also, the BMev is suggested to be a viscoelastic material in many studies (18, 21, 34, 53). To explore this further, we have characterised the viscoelastic properties by indentation creep measurements.

### Point indentation-creep measurements demonstrate the bone microenvironment can be described as a heterogeneous viscoelastic Kelvin-Voigt solid

To extract the viscoelastic properties of the BMev, an equation based on a three elements Standard Linear Solid (SLS) model (Fig. S1f) developed by Cheng *et. al*. (48) was first used to fit the indentation change (*Δδ*) *vs*. creep time (*t*) curves (Fig. 1d-e). The *Δδ*-*t* curves from 90% of the total 779 measured positions in all BMev regions of bones from both young and mature mice could be fitted well by this simple viscoelastic model (*R*^2^ > 0.9). This supports the premise that the BMev is viscoelastic rather than purely elastic.

The moduli presenting both the instantaneous elastic behaviour (*E*_*1*_) and the delayed elastic behaviour (*E*_*2*_) could be obtained from the SLS model (Fig. S3). It is notable that in the majority of cases *E*_*1*_ was significantly higher than *E*_*2*_. For all positions that were well fitted by the SLS model, 93% have *E*_1_ > 10*E*_2_ and 84% have *E*_1_ > 100*E*_2_ (Fig. S3d). This indicated any strain caused by the instantaneous elasticity was negligible. Thus, the complete SLS model is largely unnecessary and can be further simplified to a two element Kelvin-Voigt (K-V) model (Fig. S1g). The *Δδ vs*. *t* curves were then fitted to the simplified equation (for development and validation see the Supporting Material) based on the K-V model. Compared to using the SLS model, a similar number of measured positions could be fitted well (*R*^2^ > 0.9). This suggests the K-V model is sufficient to describe the viscoelasticity and the majority of BMev acts like a Kelvin-Voigt solid in nature. A K-V type solid is predominantly a viscous liquid at short time scales and predominantly an elastic solid at long time scales. This is biologically meaningful in the context of the BMev. Being a K-V type solid, ensures fast energy damping through the BMev in reaction to any abrupt mechanical loading, as well as maintaining a stable shape in the long term as part of the body scaffold.

The Young’s modulus determined by the K-V model *E*_*K-V*_ and viscosity *η* for the different bone regions are shown in Fig. 3. The median *E*_*K-V*_ for the different regions of interest in the BMev, represented in Fig 3a, are (i) cortical bone: 0.86 kPa, (ii) growth plate: 1.5 kPa, (iii) metaphysis: 0.40 kPa and (iv) bone marrow: 0.14 kPa. The mean *E*_*K-V*_ for the different regions of interest in the BMev are (i) cortical bone: 4.2 kPa, (ii) growth plate: 2.9 kPa, (iii) metaphysis: 1.7 kPa and (iv) bone marrow: 0.52 kPa. These values are significantly higher than the *E*_*H-S*_ values obtained from the pure elastic fit (as in Fig. 2a, cortical bone: *p* < 0.01, growth plate: *p* < 10^−33^, metaphysis: *p* < 10^−18^, marrow: *p* < 10^−22^), which is not surprising since the latter does not take the presence of the viscous effect into account. The *E*_*K-V*_ distributions of all BMev regions (Fig. 3c) cover several orders of magnitude, with significant differences (*p* < 0.05) found between the bone regions.

**Figure 3.**
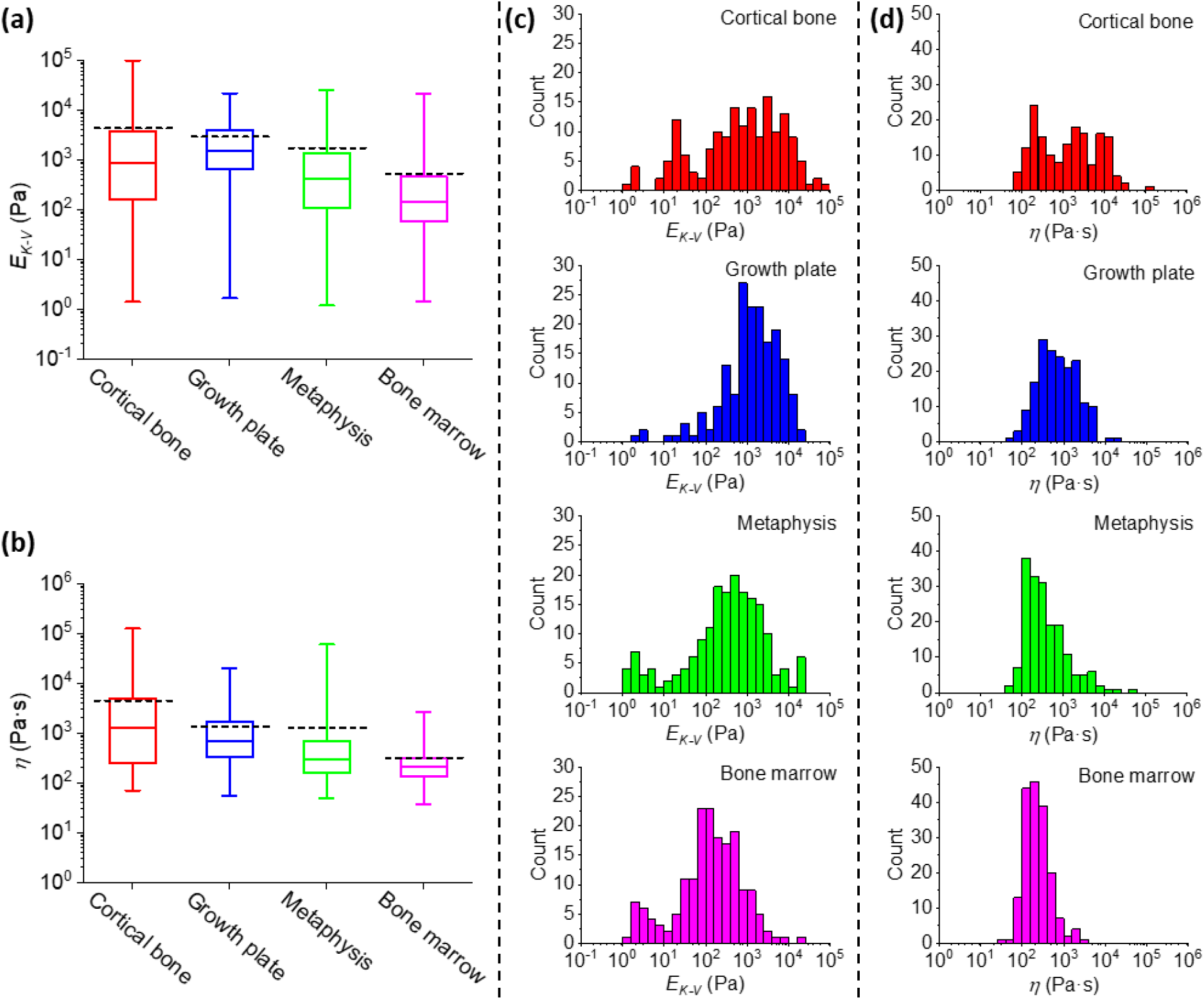
Viscoelastic properties of different bone regions (coloured version available online). (a) The Young’s modulus *E*_*K-V*_ and (b) viscosity *η* of different bone regions calculated from fits to creep curves using the viscoelastic Kelvin-Voigt model. The central box spans the lower quartile to the upper quartile of the data. The solid line inside the box shows the median and whiskers represent the lower and upper extremes. The mean values are indicated by black dashed lines. Data were obtained at the same positions as elastic measurements (Fig. 2) from all mice bones. The results are from the mean value from all repeated measurements at each position. Results from low quality fittings (*i.e*. R^2^ < 0.9) were discarded. Histograms of (c) *E*_*K-V*_ and (d) *η* of different bone regions are also shown.

The median resultant viscosity *η* of different BMev regions are represented in Fig. 3b, as (i) cortical bone: 1.3 kPa⋅s, (ii) growth plate: 0.68 kPa⋅s, (iii) metaphysis: 0.29 kPa⋅s and (iv) bone marrow: 0.21 kPa⋅s. The mean resultant viscosity *η* of different BMev regions are (i) cortical bone: 4.4 kPa⋅s, (ii) growth plate: 1.3 kPa⋅s, (iii) metaphysis: 1.3 kPa⋅s and (iv) bone marrow: 0.31 kPa⋅s. The width of *η* distribution (Fig. 3d) is smaller than that of *E*_*K-V*_, but still spans several orders of magnitude. Significant differences between different bone regions (*p* < 0.01) are also found as for the elasticity data. The creep time *τ*, defined as the time for the strain to decay to 1/*e* of its total change, can be calculated from the values of *E*_*K-V*_ and *η* of each measured position by *τ* = 3*η*/*E*_*K−V*_. The median creep time *τ* of different BMev regions are (i) cortical bone: 2.2 s, (ii) growth plate: 1.3 s, (iii) metaphysis: 2.1 s and (iv) bone marrow: 4.1 s. The mean creep time *τ* of different BMev regions are (i) cortical bone: 140 s, (ii) growth plate: 16 s, (iii) metaphysis: 34 s and (iv) bone marrow: 36 s. The median creep time for all regions is similar to the indentation time in the elastic measurements, so creep relaxation and indentation are most likely occurring simultaneously in force-distance experiments. As such, the viscous drag force is likely to have a strong effect on the *F*-*δ* curve, especially in softer areas and around the contact point, where the loading rate is highest. This is most likely the main reason for the discrepancy between the Hertz-Sneddon fit and the modulus obtained from the creep data.

It is difficult to make comparisons with the previous published studies quantifying the viscoelastic properties of the hard regions of the bone, as these studies have measured the potential load bearing mechanics (i.e. mineralised regions) (18–20) while the current study focused on obtaining measurements from the cellular microenvironment. Studies of the metaphysis have measured trabecular bone (23, 34), whereas our measurements are dominated by a more marrow-like component which accounts for most of the surface area in the region quantified in our samples.

In contrast to the hard bone regions, the composition of bone marrow in previous studies are more comparable to the values reported here. The Young’s modulus from both the elastic (*E*_*H-S*_) and viscoelastic (*E*_*K-V*_) characterisation in our study, is in good agreement with the effective Young’s modulus obtained in (39). The viscosity *η* from rheological measurements (29–31) ranges from far below 1 Pa⋅s to approximately 100 Pa⋅s. However, the resultant median and mean value of the bone marrow viscosity is significantly greater (0.2 k Pa⋅s and 0.3 k Pa⋅s) in our study. This can be explained by both sample preparation and instrument limits. The preparation and post-processing of the sample have been shown to affect the bone marrow viscosity. For instance, the viscosity of samples undergoing a freeze-thaw cycle prior to testing was lower by an order of magnitude compared to the fresh samples (31). Also, in our studies, the bone marrow was not extracted from the medullary cavity. Thus, connections of the marrow and bone tissues, and subsequently the rigidity of bone marrow structure, were maintained and relatively intact.

It is notable that viscosity is often a function of velocity and consequently varies with different measurement frequencies used in different studies. The viscosity of bone marrow has been shown to decrease as the shear rate increases according to a power law (31, 39). In contrast to dynamic measurements, the quasi-static method used in our study does not cover a broad range of frequencies. This quasi-static method requires the creep time *τ* of the sample to be significantly larger than the time required to overcome system inertia during the transition from indentation to dwelling (approximately 0.05 s). As *τ* is correlated with both *E*_*K-V*_ and *η*, the lower limit in detectable *τ* indicates that the resultant viscosity *η* using such a quasi-static method cannot be found to be much lower than the lowest values we measured (or *E*_*K-V*_ much higher than current highest values we measured).

### Aging impacts on the subcellular mechanical properties at the macro-scale of bone marrow but only minor effects on other regions of the bone microenvironment

Bone turnover in health and disease such as osteoporosis and cancer-induced bone loss show age dependent features (26, 49, 54–57). Therefore, comparing the mechanical properties of the BMev from mice at different ages will elucidate changes occurring in the mechanics of the BMev during skeletal maturation. The Young’s modulus from the elastic model, *E*_*H-S*_, together with the viscoelastic parameters *E*_*K-V*_ and *η* are classified into two groups: data obtained from young mice (age 6 to 8 weeks old, *blue*) and mature mice (age 11 to 13 weeks, *red*) (Fig. 4). Two groups of data in each region were compared by the Kruskal Wallis (KW) method.

**Figure 4.**
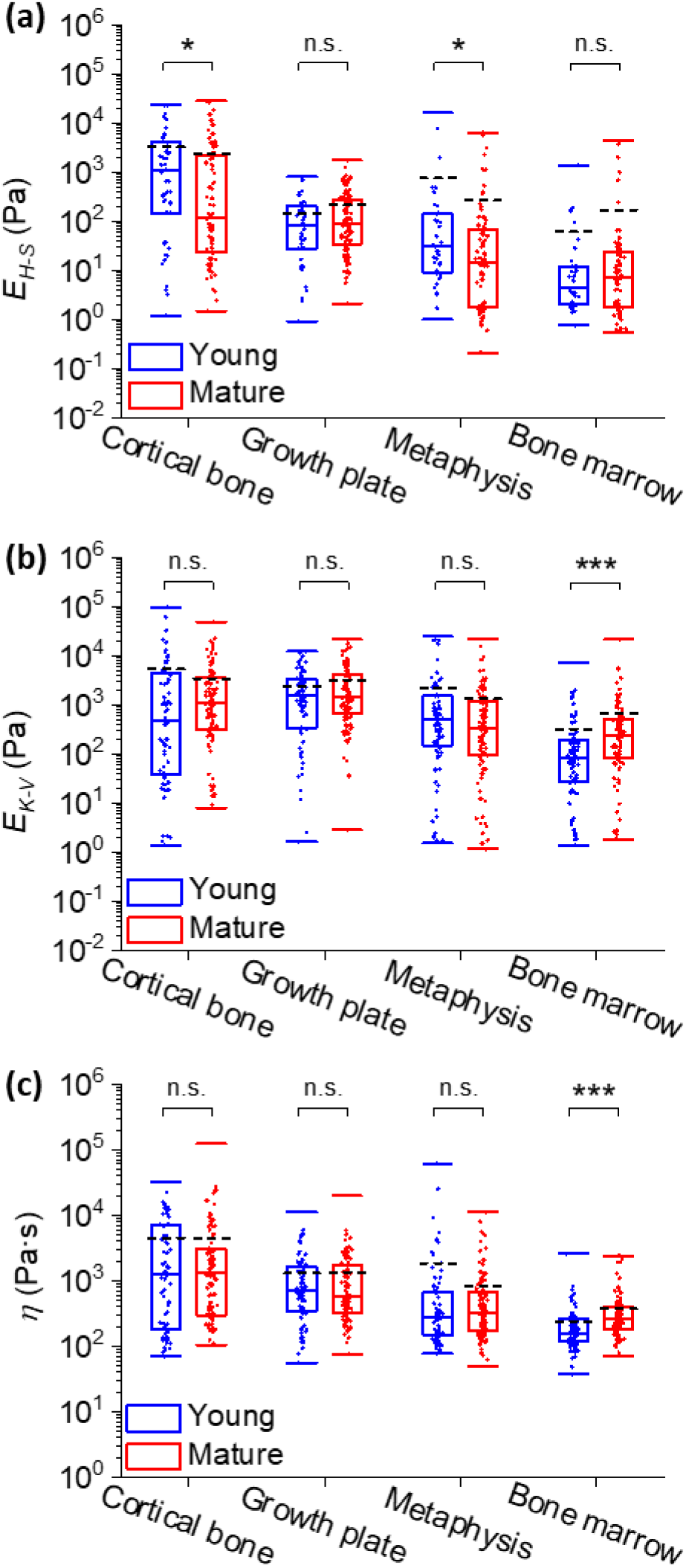
Comparison of the mechanical properties of bones from young (*blue*) and mature (*red*) mice (coloured version available online). (a) The Young’s modulus *E*_*H-S*_ obtained from the H-S model. Data are identical to those in Fig. 2. (b) The Young’s modulus *E*_*K-V*_ and (c) viscosity *η* obtained from the K-V model. Data are identical to those in Fig. 3. The central box spans the lower quartile to the upper quartile of the data. The solid line inside the box shows the median and caps below and above the box represent the lower and upper extremes. The mean values are indicated by black dashed lines. Dots represent individual data points overlaid on top of the boxplots. Data include results from both young and mature mice. The significance of statistical comparisons using the Kruskal Wallis method have been indicated above each group of boxplots (n.s.: not significant; *: *p* < 0.05; ***: *p* < 0.001).

The KW test comparing *E*_*H-S*_ (Fig. 4a) shows a minor yet statistically significant difference (0.05 > *p* > 0.03) in the cortical bone and the metaphysis. Whereas the *E*_*H-S*_ of growth plate and the bone marrow did not vary with age. For *E*_*K-V*_ and *η* (Fig. 4b-c), the KW test demonstrates a high level of significance (*p* < 0.001) in bone marrow but not for any other BMev region. There are differences in the statistical comparison of pure elastic fit results (*i.e. E*_*H-S*_) and viscoelastic fit results (*i.e. E*_*K-V*_ and *η*). The main reason is most likely that the Hertz-Sneddon model does not adequately describe the BMev properties due to not considering the viscous force, especially in soft areas such as bone marrow.

Very few studies have been performed quantifying the mechanical properties of the BMev in relation to age. Tensile tests on whole bones reported in 1976, showed only moderate differences between bones from humans at different ages (54). Recently, micro-/nano-indentation tests revealed that the mechanical properties of human cortical (55) and trabecular (56) bones were remarkably constant as a function of age. The elastic modulus of murine growth plate are reported to be significantly different between the embryonic stage and postnatal stage, age 2 week, but did not vary significantly through the next growth stages until adulthood (up to age 4 months) (25). These results are consistent with our findings. Dynamic stiffness of bone marrow stromal cells were significantly higher in adult horses compared to foals (58), which is in agreement with the age dependent trend of bone marrow mechanics observed in our study.

Such statistical comparison reflects the impact of aging on the subcellular mechanical properties of different regions of the BMev only at macro-scales. It would be helpful to integrate the impacts of aging and different length scales in future studies.

### Force mapping at the supracellular scale reveals high local mechanical heterogeneity within the different regions of the bone microenvironment

The data discussed in the previous sections were from point measurements, which does not inform on the spatial distribution of mechanical heterogeneity at the cellular scale. To explore this further, we carried out AFM force mapping. The maximum area for AFM force mapping in our setup is 100 × 100 µm^2^ with each pixel in the map down to 1 µm^2^. This ensures AFM mapping is sufficient to resolve the subcellular mechanical profile at a supracellular scale, considering the dimensions of most cells in bone (mean diameter ranging from 0.5 to approximately 100 µm) (59–61).

Fig 5a-c show representative maps obtained from cortical bone. The trigger point map (Fig. 5a) reveals the surface topography of the BMev at the trigger force. Tightly packed structures are visible (as indicated by the dashed circle), the size and shape of which are consistent with those expected for individual cells. An approximation to the surface topography of the BMev at zero load is given by the contact point map (Fig. 5b). This appears significantly smoother than the trigger point map, due to the minimal indentation at the contact point and indicates that the bone surface after splitting is relatively smooth on a supracellular scale. The lateral resolution of both maps is limited by the colloidal probe and its convolution with the surface topography. Black pixels in Fig. 5b represent missing data, due to the force distance curves at these pixels being unsuitable for fitting (see Methods). These include the large black regions indicated by arrows where the topography exceeds the lower and upper limits of the AFM’s vertical scan range. The corresponding map of *E*_*H-S*_ is shown in Fig. 5c. The *R*^2^ is greater than 0.9 for 54% and 0.5 for 95% of all curves in this map, and independent of *E*_*H-S*_ values.

**Figure 5.**
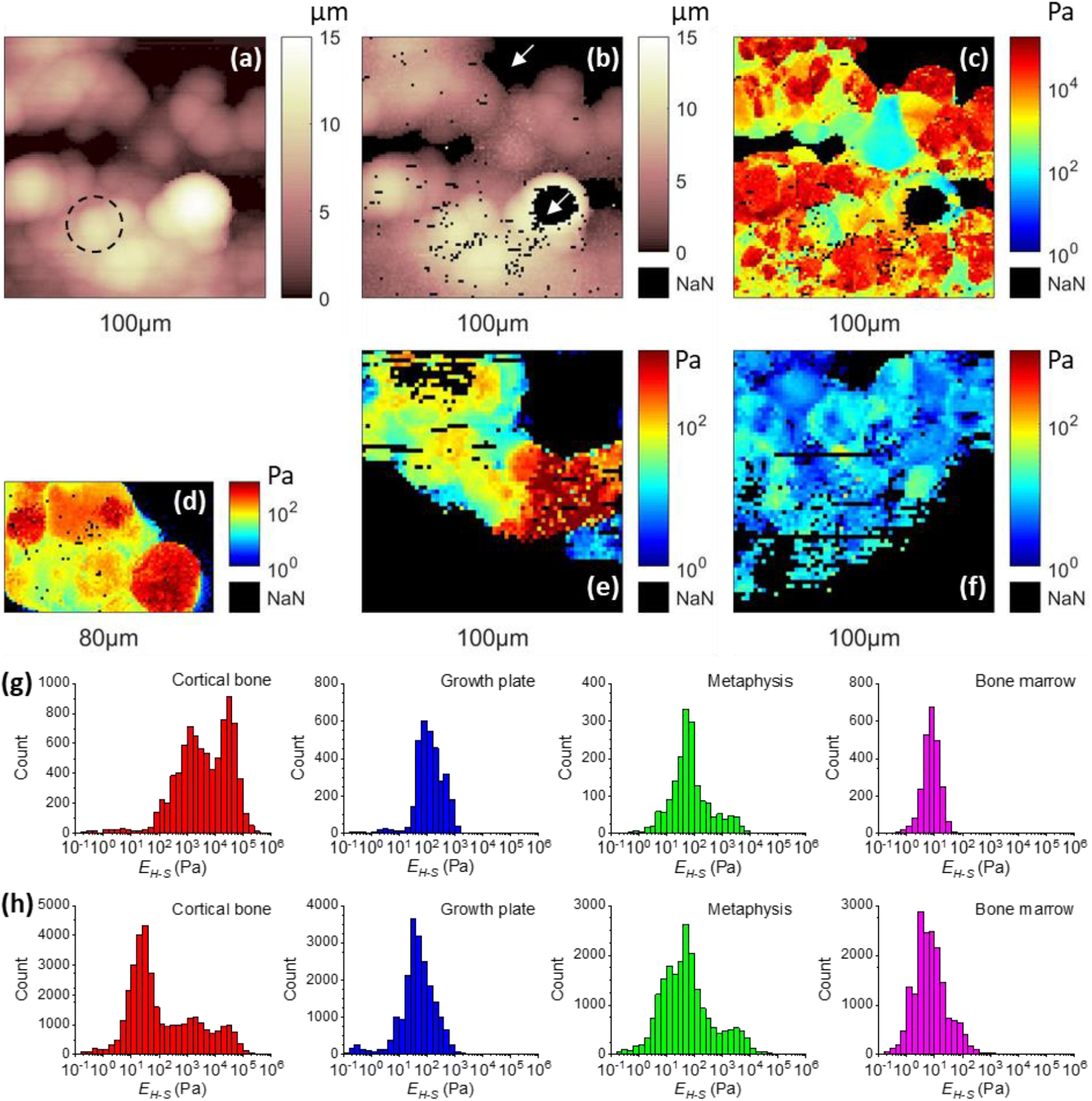
Mechanical heterogeneity of different bone regions at supracellular scale (coloured version available online). (a-c) Example of AFM maps obtained from a randomly selected position on the cortical bone. (a) Topographic map showing the height at which the trigger force was reached. Dashed circle highlights an observed spherical structure. (b) Topographic map showing the height at which the probe first makes contact with the surface. The position of the contact point was determined by the triangle thresholding method as in Fig. S1b. (c) Map of the measured Young’s modulus *E*_*H-S*_ from all force curves in the map. Curves with strongly tilted baselines (*i.e*. *d*_*b*_ > *d*_*c*_/4, as in Fig. S1c) were discarded. (d-f) Examples of *E*_*H-S*_ maps obtained from (d) growth plate, (e) metaphysis and (f) bone marrow. The data include values from curves with strongly tilted baselines (*i.e*. *d*_*b*_ > *d*_*c*_). For all maps in (a-f), missing data (*i.e*. no proper force-indentation curves could be fitted) are indicated by black pixels (by white arrows). The size scale of the maps has been indicated at the bottom of each map. (g) Histograms of the *E*_*H-S*_ distribution compiled from the maps in (c-f). (h) Histograms of the *E*_*H-S*_ distribution for different bone regions compiled from all AFM maps. For each region, at least 5 maps were recorded and each map was collected from a different mouse.

Interestingly, the mechanical heterogeneity is correlated with the topographic structures in Figs 5a and 5b, and thus helps to distinguish cellular and acellular components (the former is normally stiffer due to the nuclear, cytoskeletal and membrane components) and mineralisation status (mineralised components are expected to be much stiffer). These maps clearly show the mechanical complexity of the BMev at the supracellular scale, which is likely to be important for processes such as cell migration and proliferation that are mediated by the mechanical properties of both cells and surrounding extracellular matrix (2, 62).

Structure-correlated mechanical heterogeneity is found in all regions of interest in this study (Fig. 5d-f & S4) with *E*_*H-S*_ values covering several orders of magnitude. The structures and mechanical properties reflected in the AFM maps vary from region to region, and are difficult to accurately quantify as some areas are more difficult to map due to their large topography. Semi-quantitative comparison of the mechanical properties of different bone regions is feasible, by comparing the histograms of *E*_*H-S*_ calculated from the maps.

The histograms of *E*_*H-S*_, corresponding to the Young’s modulus maps in Fig. 5c-f, are shown in Fig. 5g. The histograms representing *E*_*H-S*_ obtained from all maps of each bone region (minimum 5 maps per region) are shown in Fig. 5h. These histograms include data from all *F*-*δ* curves with no selection criteria for curve or fit quality. As discussed in the Supporting Material, removal of poor curves has little effect on the distributions, so map data can be compared directly with individual curves from the point measurements (Fig. 2b). For all regions except the cortical bone, the shape of the distribution of *E*_*H-S*_ obtained from a single map (Fig. 5g), all maps (Fig. 5h) and individual curves covering the entire region (Fig. 2b) are comparable. The main peak in each histogram shifts by less than one order of magnitude between the different histograms for each bone region. This shows that the degree of elastic heterogeneity at the supracellular scale is comparable to the heterogeneity at a macroscopic scale. In contrast, the histograms from cortical bones are significantly different from each other. Compared to the profile of the entire cortical bone region (Fig. 2b), the peak of *E*_*H-S*_ distribution of the selected force map shifts by more than one order of magnitude. Also, the shape of the histogram for all maps obtained on the cortical bone are highly distinct from that of the histograms of either the selected map or the individual curves over the entire cortical bone region. This indicates that the overall heterogeneity of the subcellular mechanical properties in cortical bone results not only from the heterogeneity at the supracellular scale, but also variations at a macroscopic scale over the region.

It is notable that the approach speed used in force mapping was significantly increased (30 µm/s, compared to 1 µm/s in point measurements). Correspondingly, the indentation time of the majority of *F*-*δ* curves in the maps is generally not greater than 0.3 s. This is significantly shorter than the mean/median creep time *τ* of each of the BMev regions, as described in the viscoelastic section. Within such short time scales, the BMev will be predominantly viscous as a K-V type solid (see Supporting Material). We should therefore regard the modulus values obtained from indentation as an effective modulus, specific to strain rate, and encompassing both elastic and viscous contributions.

## CONCLUSION

In this work, we have used minimally processed murine bone samples to quantify the micro-mechanical properties of four distinct regions of interest using AFM. The mechanical profile of each bone region at macro-scale, revealed by point force-indentation or creep measurements using pure elastic or viscoelastic models, was found to be highly heterogeneous. The mechanical properties of different BMev regions have also shown significant differences. Moreover, aging was found to strongly impact the viscoelastic properties of the region comprising the bone marrow while having no or minimal effects in other regions of interest. All bone regions contained extremely soft areas (*i.e.* the Young’s moduli from both elastic and viscoelastic models was as low as a few Pascal), and the overall moduli were much lower than the values reported from previous studies performed on individual cells *in vitro*, whole tissues *ex vivo* or highly processed bone samples. Indeed, the viscoelastic model described the mechanical response better than the elastic model. The majority of the BMev within all regions of interest acted as a Kelvin-Voigt type solid.

This study demonstrates the feasibility of obtaining high-resolution AFM data and maps reflecting both 3D morphology and mechanical properties from complex bone tissues. AFM maps revealed that the mechanical properties of all bone regions are highly heterogeneous at the cellular/supracellular scales, ranging over 3 to 5 orders of magnitude. Such unique mechanical architecture may impact on a substantial number of active biological processes, such as bone remodelling, vascularisation, hematopoiesis and on cancer-induced bone disease. With further improvements, such as combining with post-AFM optical images, this AFM based system will be a powerful tool for further characterisation of bones in the presence of stimuli (*e.g*. hormones, cancer cells and therapeutics) or addressing mechanically relevant questions on other complex biological structures.

## SUPPORTING INFORMATION

The Supporting Material contains supplementary methods and figures S1–7.

## AUTHOR CONTRIBUTIONS

XC designed the study, performed the experiments, analysed and interpreted the data, and wrote the manuscript. RH prepared the bone samples, interpreted the data and wrote the manuscript. NM provided support for the experiments, interpreted the data and wrote the manuscript. RJH improved the theoretical model, interpreted the data and wrote the manuscript. IH, NJB and JKH designed the study, interpreted the data, wrote the manuscript and directed the project.

## ACKNOWLEDGEMENTS

This research was supported by Cancer Research UK and the Engineering and Physical Sciences Research Council (Grant Number: A21082). We thank Prof. Keith Hunter, Dr. Ashley Cadby and Miss Natasha Cowley (University of Sheffield) for fruitful discussions.

